# Semantic plasticity across timescales in the human brain

**DOI:** 10.1101/2024.02.07.579310

**Authors:** S.H. Solomon, K. Kay, A.C. Schapiro

## Abstract

Our representations of the world need to be stable enough to support general knowledge but flexible enough to incorporate new information as our environment changes. How does the human brain manage this stability-plasticity trade-off? We analyzed a large dataset in which participants viewed objects embedded in thousands of natural scenes across many fMRI sessions. Semantic item representations were located by jointly leveraging a voxelwise encoding model to find reliable item representations and a word-embedding model to evaluate semantic content. Within the medial temporal lobe, semantic item representations in hippocampal subfield CA1, parahippocampal cortex, and perirhinal cortex gradually drifted across a period of multiple months. Whole-brain analyses revealed a gradient of plasticity in the temporal lobe, with drift more evident in anterior than posterior areas. On short timescales, rapid plasticity was observed only in parahippocampal cortex, such that item co-occurrence statistics warped item representations within a single session. Together, the results suggest that the brain solves the stability-plasticity trade-off through a gradient of plasticity across semantic regions.

## 1. Introduction

The human brain flexibly adjusts its representations of the world as the environment continually changes, yet these representations remain stable enough to be useful over time. This trade-off between stability and plasticity is a pervasive tension that manifests broadly within biological and artificial systems (Meyer et al., 2014; Wandell & Smirnakis, 2009; Carpenter & Grossberg, 1987). When it comes to our units of knowledge about the world (“concepts”), this knowledge must be mutable enough to be molded by new semantic content and associations but resilient enough to not be completely overwritten. Semantic memory encompasses the multifaceted organized knowledge (e.g., perceptual, emotional, relational) that enables us to recognize, comprehend, and make inferences about objects and events (Smith & Estes, 1978; Rogers & McClelland, 2008; Binder & Desai, 2011). Semantic plasticity is evident across short and long timescales as our experiences accumulate over days and across a lifetime. For example, your concept of *crickets* likely includes that they are insects that can chirp and jump and are found among trees and patio furniture in summers. However, if you visit a street vendor in Thailand, your concept will update accordingly: crickets are also a crunchy, nutty-flavored fried snack found alongside beer and peanuts. Your *cricket* concept is malleable enough to absorb these new data without your forgetting everything you previously knew about crickets. Concepts can be substantially modified as new information is learned, as in this example, but can also undergo subtle shifts in response to a constantly changing environment (Rogers & McClelland, 2004; Musz & Thompson-Schill, 2015).

One type of semantic information that is updated in this scenario is the set of items with which crickets are associated. You previously knew that crickets co-occur with patio furniture and trees, and now you also know that they co-occur with beer and peanuts. Semantic structure—i.e., the web of associations between semantic items or features— is a fundamental component of semantic knowledge. Associations among semantic features have consequences for how categories are learned (Solomon & Schapiro, 2023) and how their labels are used in language (Solomon et al., 2019). Encoding associations between concepts themselves builds our “thematic” semantic knowledge, which includes contiguity relations based on co-occurrence in events or scenarios (Mirman et al., 2017) and influences semantic behavior such as relatedness judgments and word recall (Wisniewski & Bassok, 1999; Cann et al., 2011). If two items co-occur often, they likely play complementary roles within situations or events and are judged to be semantically related (e.g., *flower* and *vase*; *cake* and *birthday candle*). Therefore, updating this associative structure has semantic consequences: realizing that crickets can co-occur with beer and peanuts infuses the *cricket* concept with new semantic information. We do not yet know how the human brain dynamically updates representations of semantic structure within a constantly changing environment.

Semantic plasticity requires new structure to be learned, and this new information must then be integrated into existing semantic representations. Much is already known about how humans learn *novel* structure when it is isolated from our existing world knowledge. In a phenomenon known as statistical learning, humans can implicitly learn associative structure embedded among stimuli across a wide range of domains (e.g., Saffran et al., 1996; Frost et al., 2019). This learning occurs automatically, even when the structure is irrelevant to the task. The neural correlates of this learning have been thoroughly explored and suggest important contributions from the medial temporal lobe (MTL), including the hippocampus (Covington et al., 2018; Schapiro et al., 2012; 2014; 2016; Turk-Browne et al., 2009; 2010; Bornstein & Daw, 2012; Harrison et al., 2006; Strange et al., 2005). Neural representations corresponding to individual items shift as a result of structure learning, with associated items evoking more similar neural patterns post-learning (Schapiro et al., 2012, 2016; Wammes et al., 2022). Our computational model of the hippocampus implicates subfield CA1 in particular in the extraction of novel structure (Schapiro et al., 2017; Sučević & Schapiro, 2023).

Understanding how newly experienced semantic structure is incorporated into existing knowledge representations also requires an understanding of how that structure is represented in long-term memory in the first place. While the precise neural underpinnings are still unknown, temporo-parietal cortex is often implicated in language-based tasks of thematic knowledge (Mirman et al., 2017). In the context of (non-language) visual tasks, experiments have localized and examined the neural representation of object associations. In particular, the parahippocampal cortex has been implicated in visual object semantics (Bonner et al., 2016) and, more specifically, appears to represent the long-term co-occurrence statistics of objects in natural scenes (Stansbury et al., 2013; Bonner & Epstein, 2021).

This leads to our key question: How is new information integrated with these existing representations of semantic structure? More generally, the question of how new information is integrated into long-term memory is an active area of inquiry within biological and artificial systems (McClelland et al., 1995, 2020; Hebscher et al., 2019; van Kesteren et al., 2012). Systems consolidation theories posit that new declarative memories are first encoded in the hippocampus and then integrated slowly, over time and sleep, into neocortical sites (McClelland et al., 1995; Squire & Zola-Morgan, 1991; Diekelmann & Born, 2010). However, recent theories suggest that direct rapid cortical learning can occur in certain circumstances, particularly when the new information is consistent with prior knowledge (McClelland et al., 2020; Hebscher et al., 2019; van Kesteren et al., 2012). This suggests that while substantial updates to semantic knowledge may require a hippocampus-dependent gradual consolidation process, more subtle semantic shifts might be implemented rapidly and directly in cortical circuits.

A phenomenon of recent interest known as “representational drift” might provide additional insight into this problem. Not only do neural activation patterns change to accommodate new learning, but they gradually change over time even in the apparent absence of new knowledge or behavior (Rule et al., 2019; Driscoll et al., 2017, 2022; Micou & O’Leary, 2023). The mechanisms underlying representational drift and how it bears on theories of cortical function are still debated, but some have suggested that it might support the kind of continual learning described above (Driscoll et al., 2022). Regarding semantic knowledge, representational drift has been compared to the continual changes of successfully learned category representations within the layers of artificial networks (Micou & O’Leary, 2023). The phenomenon of representational drift might provide leverage into understanding semantic plasticity and how it unfolds across different timescales. To our knowledge, representational drift of semantic information has not yet been investigated in humans.

Here our goal is to examine how semantic representations shift over time and in response to changing environments (Fig. 1). Despite clear flexibility within the human semantic system (Yee & Thompson-Schill, 2016; Solomon & Thompson-Schill, 2017, 2020; Solomon et al., 2019; Musz & Thompson-Schill, 2015), neural approaches to object semantics typically make an implicit assumption of representational stability. We challenge this assumption, capitalizing on a rich fMRI dataset—in which each participant viewed thousands of visual scenes over the course of many months—to explore whether and how semantic representations change over short and long timescales in the human brain. We focus on the MTL because of its known contributions to rapid learning and visual object semantics, and its theorized joint role in perceptual and memory processes (Graham et al., 2010; Martin & Barense, 2023). We uncover reliable object representations across the MTL and find that object representations in hippocampal subfield CA1, parahippocampal cortex (PHC), and perirhinal cortex (PRC) contain semantic information. Within these semantic regions, we find that object representations drift across the span of ∼8-months, with PHC showing the highest rate of drift. Finally, we find evidence that changes in these PHC object representations are explained by changes in semantic structure within the recent (∼1 hour) visual environment. Whole-brain analyses further reveal a distribution of representational flexibility across ventral temporal cortex. Together, the results demonstrate that PHC contains object semantics that dynamically update, whereas more posterior areas represent object semantics in a more stable manner. The semantic system may thus solve the stability plasticity trade-off by allowing only a subset of its representations to rapidly adapt to a changing environment.

**Figure 1:**
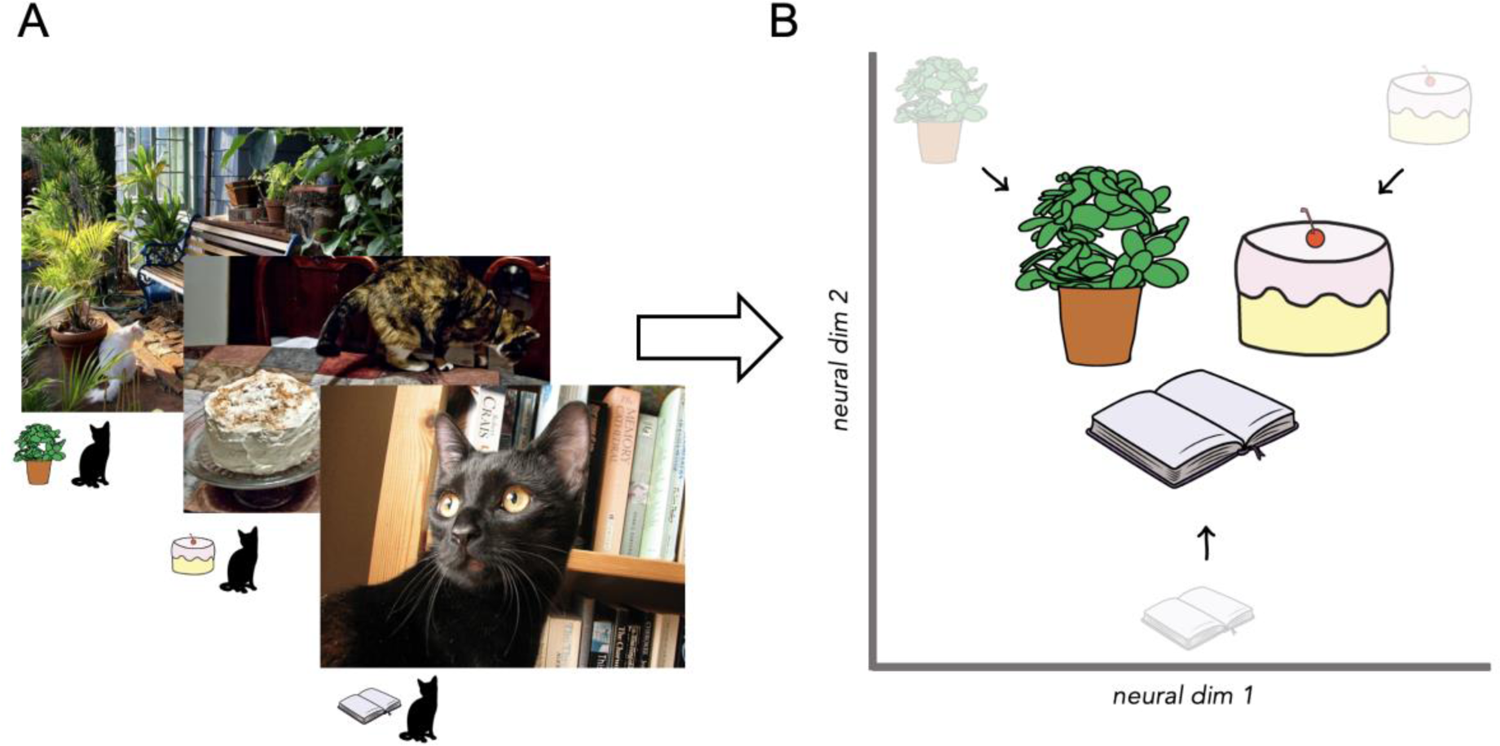
Natural scenes and item co-occurrences. (A) Stimuli consisted of natural scene photographs from the COCO image database (http://cocodataset.org). The first example image contains a cat and plants, the second contains a cat and a cake, and the third contains a cat and books. (B) We test whether items that share recent patterns of co-occurrence *(plant, cake*, and *book* all co-occur with cat) become more similarly represented in the brain. Axes are hypothetical dimensions of neural activity, and proximity of items within this space reflects the similarity of their neural representations.

## 2. Results

We analyzed data from the NSD experiment (Allen et al., 2022), in which participants viewed thousands of natural scenes across ≥30 fMRI sessions over the span of ∼8 months. The stimuli were real-world photographs containing familiar objects, including tools, vehicles, and animals (e.g., cake, plant, book; Fig. 1A). Our key goal was to explore the plasticity of semantic representations in the brain. To do this, we first implemented a voxel-wise object encoding model in order to locate reliable item (object) representations across the brain (Fig. 2A). We then found the regions in which these item representations were semantic in nature, using a word-embedding model that captures semantic similarities using word co-occurrences in text (Fig. 2B). Once semantic item representations were located, we could then explore the plasticity of these representations on different timescales. We assessed long-term plasticity by examining the extent to which item representations drifted across the course of the experiment (∼8 months; Fig. 2C). To assess short-term plasticity, we asked whether item co-occurrence statistics in the first half of a session predicted item similarities in the second half of the session (Fig. 2D).

**Figure 2:**
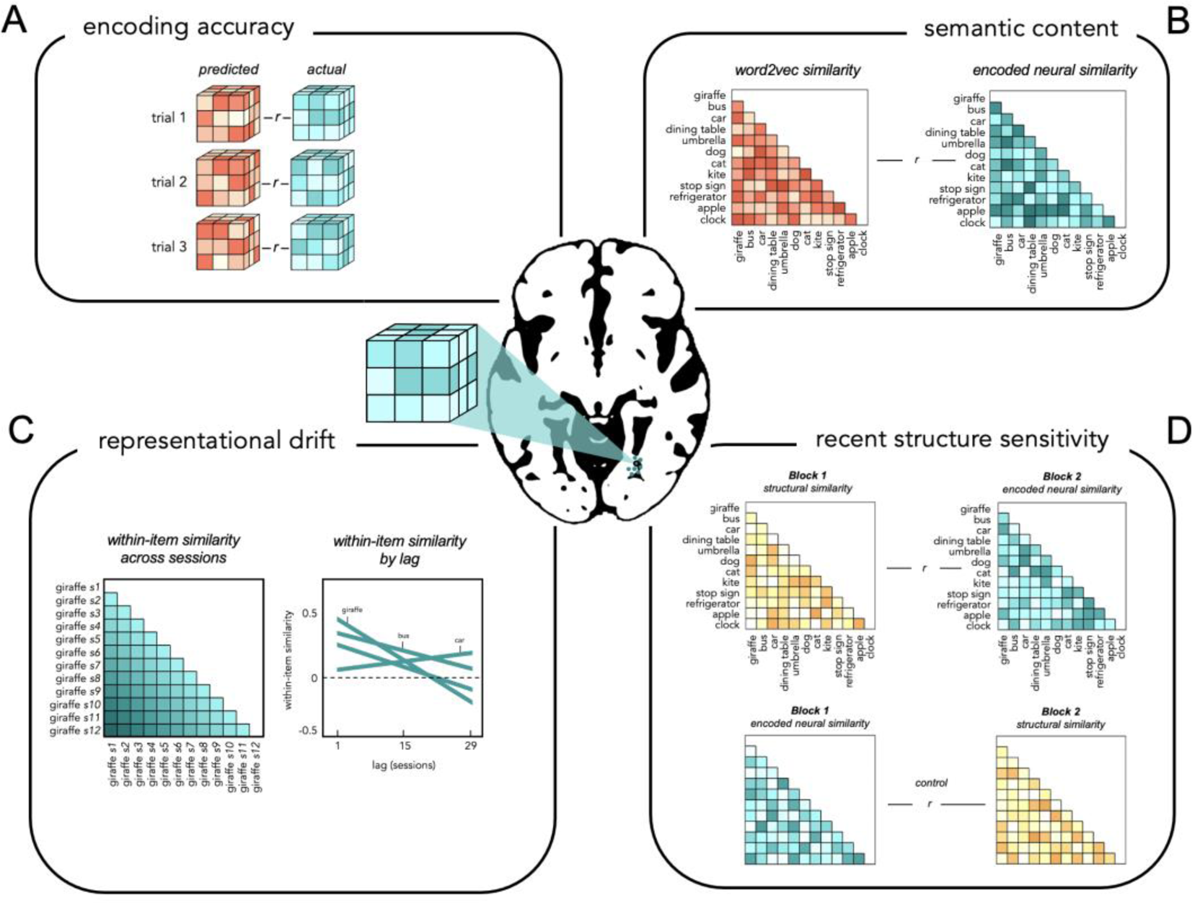
Methodological approach. Multiple measures were extracted from searchlights across the brain. (A) The fidelity of item representations was assessed using a voxel-wise encoding model that predicted each voxel’s response to each of the 80 objects. (B) Semantic content was assessed by comparing neural across-item similarity with the similarities contained within a word embedding model. (C) Representational drift corresponded to decreases in within-item neural similarity over time. (D) We calculated sensitivity to recent statistical structure of the stimuli by determining whether the statistical structure in the first half of a session influenced neural across-item similarity in the second half of a session. We subtracted the correlation in the opposite direction as a tight control to ensure our measure reflected a time-dependent influence of recent structure on neural representation.

The degree of successful item decoding, semantic content, representational drift, and sensitivity to recent statistics was measured in searchlights across the brain (Fig. 2).

Searchlight values were averaged within ROIs that included hippocampal subfield CA1; a combined ROI including hippocampal subfields CA2, CA3, and dentate gyrus (DG); the subiculum; parahippocampal cortex (PHC), perirhinal cortex (PRC), and control region V1. Note that we use the term PHC in the context of its definition within the MTL (Hindy & Turk-Browne, 2016). Whole-brain results are also reported.

### Object Encoding

We first determined which brain regions contained reliable item representations, since these item representations were the focus of our subsequent analyses. A voxel-wise object encoding model was used to determine which regions of the MTL encoded the identity of the 80 items (Figs. 2A; 3A). Successful object encoding (all *p*’s *<* 0.008) was observed in all MTL ROIs as well as in V1 (Fig. 3B). These results revealed that each of these brain regions—including the hippocampal subfields— contained reliable object representations evoked by the natural scenes. A whole-brain analysis revealed widespread encoding success across much of the brain, especially in visually responsive cortex (Fig. 3C).

**Figure 3:**
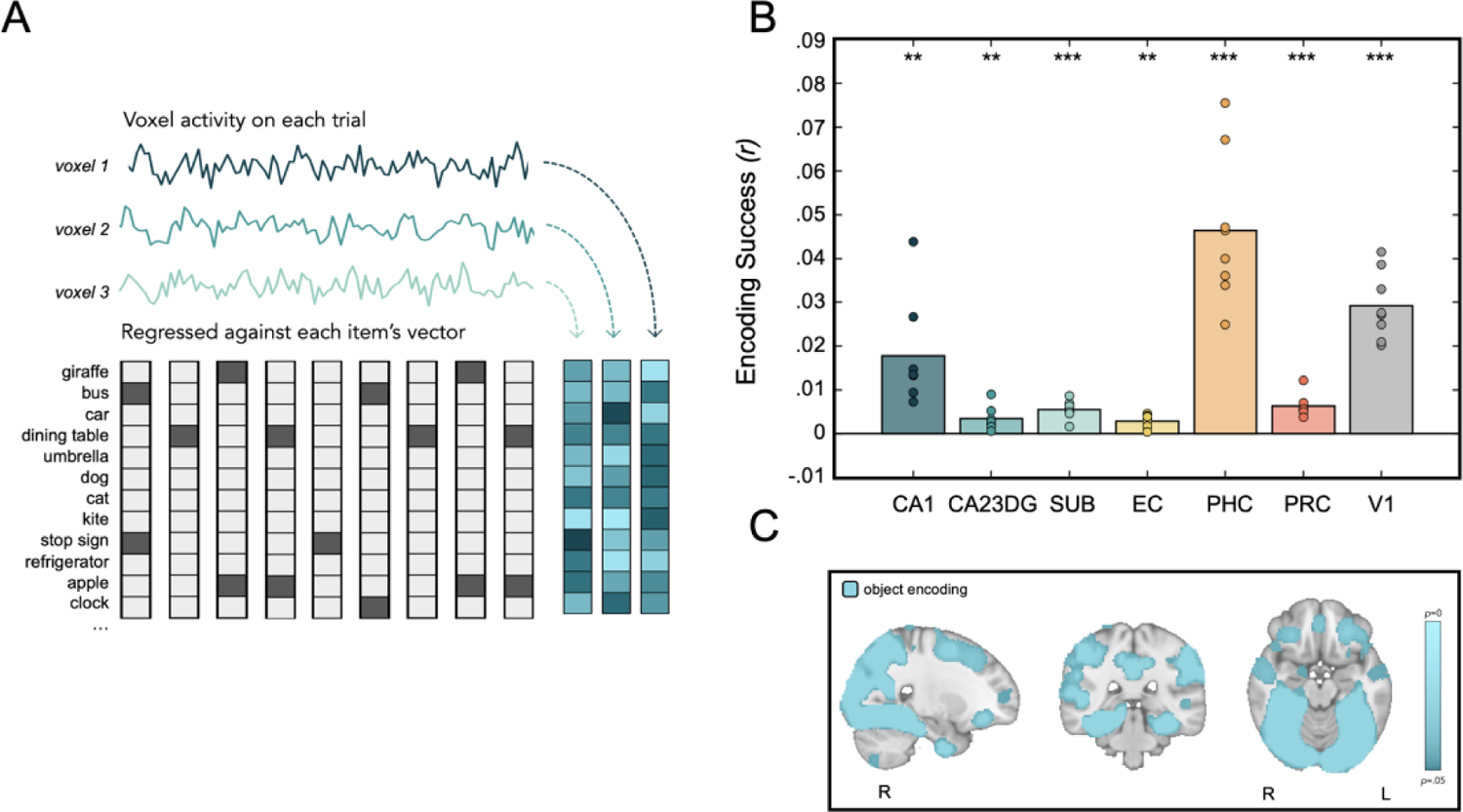
Item encoding. (A) A voxel-wise encoding model employing cross-validated ridge regression was used to predict each voxel’s response to the 80 items and to subsequently predict multivoxel activity patterns evoked by individual scenes. Predictions were compared to actual neural responses to determine the fidelity of item representations across the brain. Item encoding was successful within all ROIs (B) and large swaths of cortex (C).

### Semantic Content in the MTL

Having established the locations of reliable item representations, we then sought to determine whether these representations contained semantic information. The encoding model generated predicted patterns of neural activity for each item; these patterns were correlated with each other to generate a neural similarity matrix. Neural similarity was then compared with semantic similarity, which was captured using word2vec (Fig. 2B), a measure of similarity derived from word co-occurrences in text. Semantic similarity reliably correlated with neural similarity in hippocampal subfield CA1 (*t*(7)=3.2, *p*=0.015), PHC (*t*(7)=5.52, *p*=0.0009), and PRC (*t*(7)=2.65, *p*=0.033). Semantic similarity did not reliably predict neural item similarity in the other hippocampal subfields (CA23DG: *t*(7)=2.14, *p*=0.069; SUB: *t*(7)=2.3, *p*=0.055) or in EC (*t*(7)=1.53, *p*=0.17). A significant negative effect was observed in V1 (*t*(7)=-4.97, *p*=0.002). This negative effect can be explained by a constraint we implemented in which only pairs of items that never co-occurred were analyzed (see Supplemental Figure 1). This was done to eliminate the possibility that items appearing in the same images would inflate the similarity between neural and semantic spaces. The observed negative relationship between semantic similarity and neural similarity in V1 was likely driven by the fact that items with high semantic similarity often occur in visually dissimilar scenes—for example, snowboards and surfboards are semantically similar but occur in very different visual contexts. Thus, our results reveal that evoked object representations in CA1, PHC, and PRC are semantic in nature (Fig. 4A). A whole-brain analysis additionally revealed reliable semantic content within some of the more anterior areas with reliable item representations (Fig. 4B). Establishing semantic content within CA1, PHC, and PRC set the stage to then examine the plasticity of these semantic representations over time.

**Figure 4.**
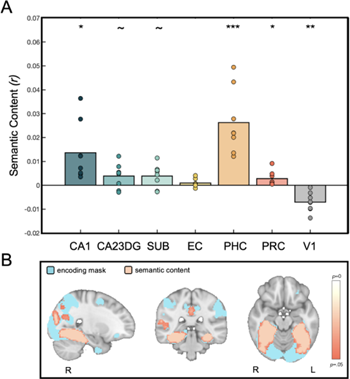
Semantic content. (A) Neural across-item similarities positively correlated with semantic similarities extracted from a word embedding model in CA1, PHC, and PRC. (B) Item representations were semantic in nature within a subset of temporal, parietal, and cingulate regions that exhibited successful item encoding.

### Long-term Drift of Item Representations in the MTL

Having established semantic representations in CA1, PHC, and PRC, we aimed to determine whether these representations gradually drifted over time. If neural activity for a given item drifts over the span of months, then within-item pattern similarity would be more similar for sessions closer together in time (e.g., sessions 1 and 2) than for sessions more separated in time (e.g., sessions 1 and 30), with the degree of similarity scaling with the number of intervening sessions (Fig. 2C). That is, within-item similarity would decrease as the number of intervening sessions increased (Fig. 5A). On average, there were approximately 8 days in between sessions. Reliable drift was observed in all semantic ROIs (Fig. 5B), including CA1 (*t*(7)=2.81, *p*=0.026), PHC (*t*(7)=3.44, *p*=0.012), and PRC (*t*(7)=3.68, *p*=0.008). Drift was not observed in V1 (*t*(7)=1.46, *p*=0.18). Drift in PHC was reliably greater than in PRC (*t*(7)=2.86, *p*=0.024) and marginally greater than in CA1 (*t*(7)=2.30, p=0.055) and V1 (*t*(7)=2.28, *p*=0.056). No other comparisons were significant (*p*’s > 0.1). This drift effect cannot be explained by an increase or decrease in semantic content within neural representations across the course of the experiment; correspondence between neural and semantic similarity spaces, established using word2vec, was stable over time, with no increase or decrease across the thirty sessions (CA1, PHC, PRC, and V1 *p*’s > 0.4). A whole-brain analysis revealed reliable drift of item representations in anterior ventral temporal cortex, predominantly in the left hemisphere. It is notable that not all regions containing semantic representations revealed the same amount of drift (Fig. 5C), suggesting a gradient of stability to plasticity within the semantic system (see Supplemental Figure 2 for further analysis). Taken together, these analyses confirm the existence of drift in semantic object representations over long time-scales and suggest that these representations may drift at different rates in different brain regions. More generally, these results demonstrate the plasticity of neural object representations, leaving open the question of what factors contribute to these representational changes.

**Figure 5.**
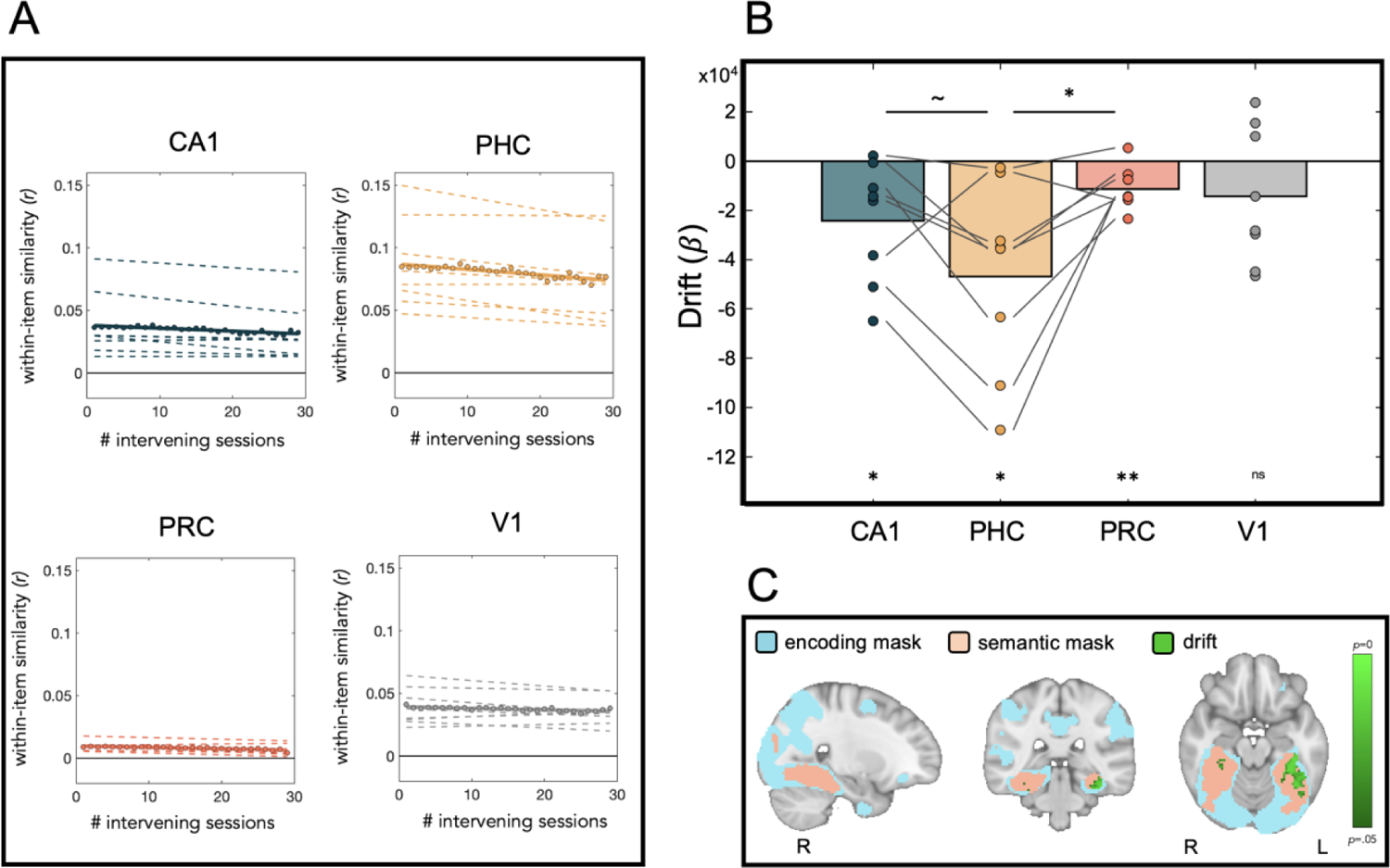
Representational drift across long timescales. Within regions containing semantic item representations, we asked whether these representations drifted across the 30 sessions (∼8 months). (A) Within-item neural similarity was regressed against number of intervening sessions. Negative slopes indicate representational drift, where item representations gradually changed over time. Circles reflect average item similarity across participants at each time interval, and the solid line indicates the linear fit. Dashed lines indicate fits for individual participants. (B) Drift was significant in all semantic ROIs but not in V1. (C) Only a subset of semantic regions exhibited significant drift in item representations over time.

### Recent Structure Rapidly Influences Item Representations in PHC

We observed that regions within the MTL (CA1, PHC, and PRC) and temporal cortex contained semantic item representations that drifted over long periods of time. Here we ask whether and where semantic shifts might occur more rapidly (within a single fMRI session), and whether the recent semantic environment can drive such shifts. To do this, we compared item co-occurrence structure in the first half of a session with across-item neural similarity in the second half of that session to determine whether recent structure in the environment results in rapid representational change (Figs. 2D; 6A). That is, if two items tend to occur with the same items in the first half of the session, are they represented more similarly in the second half of the session? Importantly, we implemented a tight control, in which we also calculated the correlation between item statistics in the second half of the session with neural similarity in the first half of a session. We subtracted this backwards correlation from the forwards correlation to ensure that our results signify a time-dependent effect of recent co-occurrence structure on neural similarity. We found that recent structure rapidly influenced the neural similarity of items in PHC (*t*(7)=3.1, *p*=0.017). This result was marginal in CA1 (*t*(7)=2.28, *p*=0.056), and was not significant in either PRC (*t*(7)=1.11, *p*=0.3) or in V1 (*t*(7)=0.42, *p*=0.68). These results suggest that the semantic plasticity in PHC is in part explained by the rapid influence of recent structure on neural object representations (Fig. 6B).

**Figure 6.**
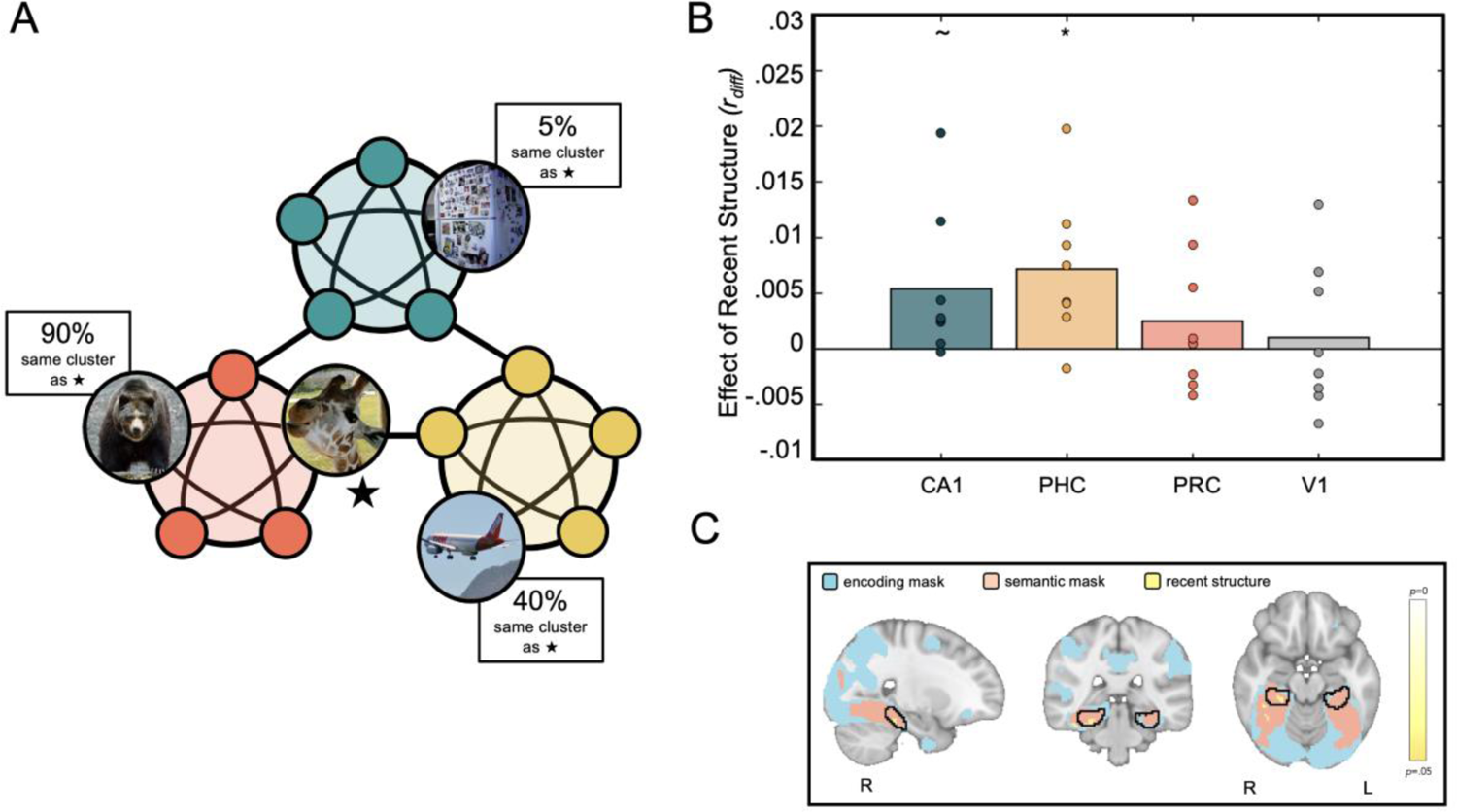
Influence of recent statistical structure on item representations. (A) Structural similarity between pairs of items was calculated using a network modularity approach. Items that tended to occur with the same items across trials have a higher likelihood of being assigned to the same module and thus were structurally “similar”. (B) For a pair of items, structural similarity in the first half of a session predicted neural similarity in the second half of a session in PHC, but not in other semantic ROIs or in V1. (C) A whole-brain analysis revealed a significant cluster that overlaps with the PHC ROI (black outline), as well as clusters in right fusiform cortex and bilateral clusters in posterior middle temporal gyrus (not shown).

We correlated the rapid structure effect in each session with a measure capturing the degree of similarity (“structure match”) between recent and long-term item co-occurrence structure. We observed that PHC representations were more likely to change when the recent structure (i.e., structure in the first half of the session) differed from long-term structure (*t*(7)=3.0, *p*=0.012). In other words, PHC updated its item representations more when it was confronted with more novel structural information (Fig. 7).

**Figure 7.**
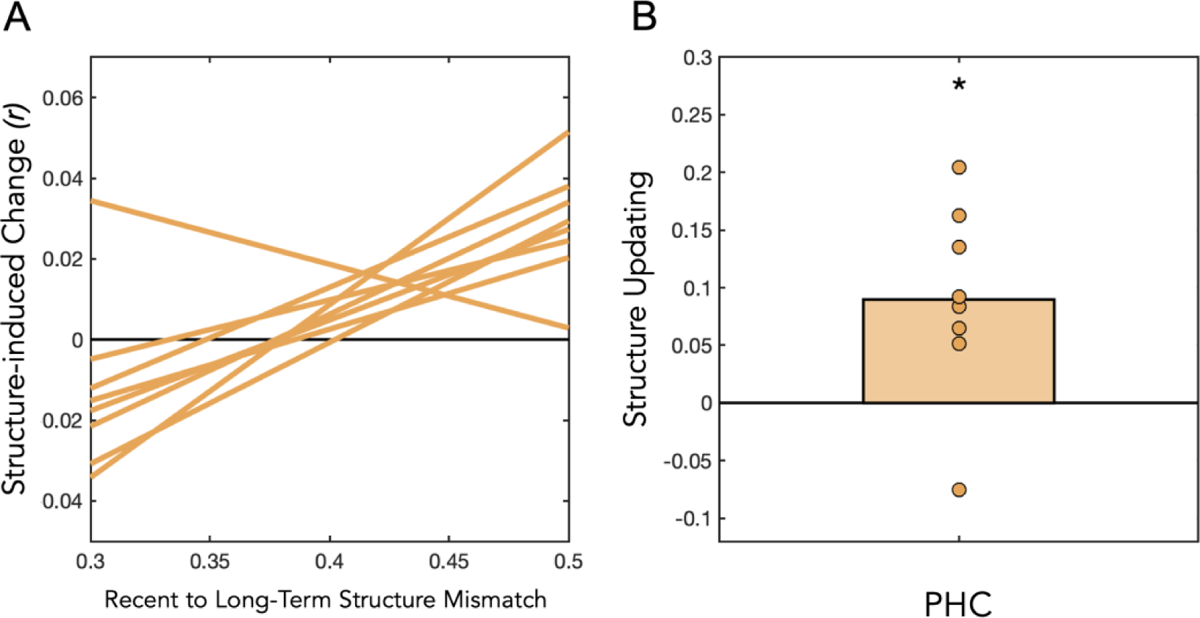
Relationship between recent and long-term structure. The similarity between long-term item co-occurrence structure and the structure in the first half of each session predicted the degree of structure-induced change in PHC, indicating that PHC updated its representations in the face of new information.

In a whole-brain analysis, we explored whether semantic regions outside the MTL also contained item representations that were rapidly influenced by recent structure. Notably, this analysis revealed a significant cluster that overlapped with our PHC ROI (Fig. 6C). Additionally, we observed a significant cluster extending into the right fusiform cortex and bilateral clusters in posterior middle temporal gyrus. The locations of the ventral temporal clusters fell anterior to the parahippocampal place area (PPA) and fusiform face area (FFA), as assessed on the cortical surface (Supplemental Figure 3).

## 3. Discussion

How plastic are semantic representations in the human brain? To address this question, we analyzed data from an fMRI experiment in which participants viewed objects embedded in thousands of natural scenes across the span of approximately eight months. We found reliable representations of the objects in large swaths of visual and temporal cortices, including all regions of interest within the medial temporal lobe (MTL). A subset of these regions contained item representations that were semantic in nature, including hippocampal subfield CA1, parahippocampal cortex (PHC), and perirhinal cortex (PRC). These semantic item representations gradually drifted across the eight months, with the highest level of drift observed in PHC. On short timescales (within ∼1 hour), representational change in PHC was predicted by the object co-occurrence structure embedded within recently observed visual scenes, suggesting that item representations in PHC are rapidly influenced by the statistical structure of the semantic environment.

Our finding that item representations in PHC are influenced by recent object co-occurrence statistics suggests that this region is involved in rapid semantic learning, at least in the context of viewing natural scenes (Fig. 1). The location of this effect is anterior to the parahippocampal place area (PPA), which is functionally defined as a region recruited during scene perception (Sun et al., 2021). The location of our learning effect relative to typical perception effects is consistent with the idea of a functional gradient in the ventral stream, in which posterior portions are more involved in visual encoding and spatial associations, whereas anterior portions are more involved in visual memory, scene context, and non-spatial associations (Aminoff et al., 2007; Bainbridge et al., 2021; Baldassano et al., 2013; Steel et al., 2023). Additionally, our observation that PHC representations are influenced by recent object co-occurrence statistics is consistent with and builds on prior work claiming that this region represents the long-term object co-occurrence structure in natural scenes (Bonner & Epstein, 2021; Stansbury et al., 2013). Here, we show that these structural representations are dynamic—they are constantly modified by recent experience.

The flexibility of item representations in PHC—evidenced by both rapid structural shifts and long-term representational drift—is an important step towards understanding how semantic plasticity unfolds within the human brain. However, PHC’s role in semantic updating more generally is unclear. It is possible that we observed representational change in PHC because its internal computations render it uniquely suited to associative contextual processing (Aminoff et al., 2007; 2013), but it is also possible that PHC emerged because of its role in the visual processing of naturalistic scenes (Epstein & Baker, 2019). It will therefore be interesting to test whether our plasticity findings hold in experiments using non-scene stimuli. PHC does seem sensitive to an object’s associations and contexts when items are shown in isolation in the absence of a scene (Aminoff et al., 2007; Bonner & Epstein, 2021). Further, neuroimaging investigations of thematic semantic processing—involving verbal rather than visual stimuli—implicate regions of posterior temporal cortex in the processing of semantic relations (Mirman et al. 2017), raising the possibility of a general role of semantic plasticity in PHC. Another possibility is that subtle representational shifts, like the ones reported here, will be found in the diverse regions of the brain responsible for representing the stimulus content itself. In our data, the item co-occurrence statistics are implicitly represented in the similarity space of the neural item representations themselves. It is thus possible that, for any given experiment, semantic plasticity may be observed within the neural regions that represent the particular stimuli used. In other words, it is not yet known whether the plastic representations observed in PHC arise by means of domain-specific or general mechanisms.

We observed only marginal evidence for an influence of statistical structure on item representations in the hippocampus. The hippocampus is known to be sensitive to statistical structure (Covington et al., 2018; Schapiro et al., 2012; 2014; 2016; Turk-Browne et al., 2009; 2010; Bornstein & Daw, 2012; Harrison et al., 2006; Strange et al., 2005; Schlichting et al., 2015), and we have argued that the properties of hippocampal subfield CA1 in particular are well-suited to rapid learning of statistical associations between stimuli (Schapiro et al., 2017). The hippocampus can learn novel associations among unfamiliar stimuli such as novel objects and fractals (Schlichting et al., 2015; Schapiro et al., 2016) but can also learn novel structure among real-world objects and/or visual scenes (Tompary & Davachi, 2017; Bornstein & Daw, 2012). However, in these paradigms, while the stimuli may either be novel or drawn from prior knowledge, the *associations* between stimuli are novel. This is in contrast to the current study, in which the real-world items within the natural scenes were embedded within real-world co-occurrence structure. That is, the statistical structure of the items was consistent with prior structural knowledge (as these were photographs from the real world; Fig. 1A). Thus, while the hippocampus is necessary for novel structure learning, it might not be necessary for making tweaks to pre-existing structural representations of known items. However, the marginal effect of rapid statistical learning in CA1 leaves open the possibility that the hippocampus did contribute to the rapid learning effects in cortical regions. Future research leveraging hippocampal lesion or perturbation is needed to determine whether the hippocampus plays a causal role.

We did not find reliable evidence of rapid learning in the hippocampus, instead observing robust neocortical learning in PHC. This is inconsistent with a strict conception of systems consolidation in which new information must first be encoded in the hippocampus and then subsequently communicated to neocortex through a gradual consolidation process (Squire & Zola-Morgan, 1991). It is, however, consistent with a growing body of empirical and theoretical work suggesting that rapid cortical learning can occur under certain circumstances. For example, rapid formation of cortical engrams and “tags” have been demonstrated during learning in rodents (Kitamura et al., 2017; Lesburguéres et al., 2011), visual exposure to movies of natural scenes results in an immediate memory trace in cat visual cortex (Yao et al., 2007), and there is MRI evidence in humans for rapid learning-related microstructural changes in parietal cortex and medial temporal lobe cortex (Brodt et al., 2018; Sagi et al., 2012). Fast mapping is another phenomenon in which incoming information putatively bypasses traditional hippocampal circuits to be integrated within neocortex (e.g., Merhav et al., 2015). By examining the conditions that support rapid cortical learning, it appears that one important factor is the extent to which new information is congruent with prior knowledge or existing “schemas” (Tse et al., 2007; Hebscher et al., 2019; van Kesteren et al., 2012; McClelland et al., 2020). When new information is consistent with what is already known, it can potentially bypass the hippocampally-mediated consolidation process and become rapidly encoded within neocortex. A recent artificial network model offers a computational demonstration of how new information that is congruent with prior knowledge can be rapidly integrated within cortex-like representations, while incongruent information requires a gradual consolidation process involving hippocampal replay of old and new memories (McClelland et al., 2020).

We observed increased rapid learning in PHC when the structure of the recent environment was more dissimilar from long-term structure, suggesting that PHC updates its representations when confronted with new information. Based on the theory that rapid cortical learning can occur only when the new information is consistent with prior knowledge, one might have predicted the opposite—that increased learning would occur when the recent and long-term structure are more similar. However, the semantic structural changes we explore here are quite subtle. Even the most “dissimilar” co-occurrence structures in our case are likely still highly consistent with prior knowledge; The stimuli were natural images of real-world scenes, with associative structure thus drawn from the known distribution of real-world statistics. Our results are therefore compatible with the view that rapid cortical learning can only occur when the new information is consistent with prior knowledge.

In addition to rapid cortical learning, we also observed representational drift of semantic content over a period of many months. To our knowledge, this is the first such demonstration of semantic representational drift in humans. The phenomenon of representational drift is typically explored in animal models. In one example, mice were trained on a virtual-navigation task such that their behavioral performance was stable across a month. Despite this stable behavior, the individual neurons in posterior parietal cortex that were sensitive to different features of the task changed across days, resulting in the drift of activity patterns over time (Driscoll et al., 2017). Representational drift can be driven by both time and experience (Geva et al., 2023), may contribute to the delicate balance between stability and flexibility in learning and memory systems (Driscoll et al., 2017, 2022), and could play a role in continual learning (Driscoll et al., 2022; Micou & O’Leary, 2023). Notably, there is recent evidence of representational drift in humans: Roth & Merriam (2023) recently used the Natural Scenes Dataset to reveal drift of representations in early visual cortex. We did not observe drift in V1 in the current study, but we targeted object representations, which would be expected to manifest more anteriorly, whereas Roth & Merriam directly targeted representations characteristic of V1 (orientation and spatial frequency).

Our joint observations that recent item statistics rapidly influenced item representations in PHC and that these representations drifted over longer periods of time leads us to speculate that representational change on these two timescales may be linked. That is, the accumulation of small representational changes based on environmental statistics may, over time, result in the gradual drift of these representations over longer timescales. The participants in the current study were regularly exposed to most of the target items (e.g., plant, book) in their day-to-day life in between fMRI sessions, providing frequent opportunities for their representations to update based on which items were seen together in different contexts. Thus, the long-term representational drift we observed in PHC would presumably mainly reflect observations of and experiences with these items and their surroundings in the real world. Future work may be able to more directly link these short- and long-term changes through exposure to stimuli unlikely to be encountered between sessions.

We found different levels of drift over time across semantic regions: even though drift was observed in PHC, CA1, and PRC, the degree of drift varied across regions, with greater drift in PHC than PRC. A whole-brain analysis suggested increased long-term plasticity in more anterior portions of semantic-sensitive regions (Fig. 5C). The finding that some regions revealed drift of semantic representations whereas others did not provides a hypothesis for the brain’s solution to the stability-plasticity trade-off in the semantic system. The brain harbors less volatile item representations in more posterior areas that are able to withstand a dynamic environment, in addition to more malleable item representations more anteriorly that can accommodate new information when needed. Additionally, only a subset of semantic regions is sensitive to the recent statistics of the environment, including PHC, further suggesting a division of labor within the brain such that some regions retain stable representations of the world whereas other regions shift their representations over short and long timescales.

In summary, our findings reveal how the brain manages the stability-plasticity trade-off in the context of visual object semantics. While some regions of visual and temporal cortex contain static representations of the world, other more anterior regions, especially PHC, contain plastic representations that absorb the recent statistics of the visual environment and also drift across long periods of time. This variation in representational flexibility may allow the brain to provide the stability that is necessary for a useful model of the world while also being able to adjust to a constantly changing environment.

## 4. Methods

This study is based on the Natural Scenes Dataset (NSD) which has been described in Allen (2022) and is openly available (http://naturalscenesdataset.org). Here, we summarize the main characteristics of the dataset and describe the data that are relevant to the current paper. See Allen et al. (2022) for full details on MRI acquisition.

### Participants

Eight participants contributed data to the study (six females and two males; age range 19-32). All participants had normal or corrected-to-normal vision. Informed consent was obtained in accordance with the University of Minnesota Institutional Review Board.

### Design and Procedure

Each participant viewed 9,000–10,000 distinct natural images across 30-40 sessions while fMRI data were collected. Each of these images was presented up to three times across the entire experiment; participants performed a long-term continuous recognition task on each trial in which they indicated via a button press whether each image presented was new or old (i.e., whether that image had previously appeared at any earlier time in the experiment). Each NSD session contained 12 runs of the NSD task, with 62-63 image trials per run, resulting in 750 image trials per session. On each trial an image was presented for three seconds followed by a one second gap. In this study, data from sessions 1-30 were analyzed for each participant, corresponding to a time range of 210-287 days.

### Stimuli

Natural images were drawn from the Common Objects in Context (COCO; https://cocodataset.org) database. COCO images are photographs that come with segmentations indicating the presence and location of a set of 80 objects (e.g., person, giraffe, bus, umbrella). For the purposes of this study, we represent each image as a sparse vector indicating the count of each of the 80 objects contained therein. Each of the 80 COCO objects appeared in at least 100 images to each participant, and 90% of the images contained multiple object categories. In this paper we use the terms “objects” and “items” interchangeably. The experiment was designed such that each participant was assigned a set of 10,000 images, 9,000 of which were unique to that subject and 1,000 of which were common across subjects; thus, different subjects experienced different object co-occurrence statistics over time.

### MRI Acquisition

The NSD neuroimaging data were collected at the Center for Magnetic Resonance Research at the University of Minnesota. Anatomical data were collected using a 3T Siemens Prisma scanner and a 32-channel RF head coil and included T1 (0.8-mm isotropic resolution, TR=2400 ms, TE=2.22ms, TI=1000 ms, flip angle 8°) and T2 (0.8-mm isotropic resolution, TR=3200 ms, TE=563 ms, TI=1000 ms) scans. Functional data were collected using a 7T Siemens Magnetom, a 32-channel-receive RF head coil, and gradient-echo EPI at 1.8-mm isotropic resolution and 1.6-s TR with whole-brain coverage. Participants were scanned approximately once per week.

### MRI Pre-processing and GLM Analysis

Data were processed using a variety of methods and multiple versions of data were generated. Here we summarize the methods that are relevant to the current study. Functional data were pre-processed to correct for slice time differences, head motion within and across scan sessions, EPI distortion, and gradient non-linearities. These data were upsampled to a 1.0-mm high-resolution preparation. GLM analysis of these pre-processed timeseries data provided beta estimates of BOLD response amplitudes for single trials (i.e., single image presentations). We used the version of the beta estimates (‘betas_fithrf’) in which an optimal HRF is identified for each individual voxel before fitting a single-trial design matrix to the data. These high-resolution 1.0-mm betas were mapped to 2.0-mm MNI space using cubic interpolation. Trial-wise activity was z-scored for each voxel in each session. To define regions of interest (ROIs) we used MTL atlas labels derived from a database of manual anatomical segmentations (Hindy & Turk-Browne, 2016; Aly & Turk-Browne, 2016).

### Object Encoding Model

An encoding model was used to localize brain regions containing reliable object representations, and to generate multivoxel object pattern estimates in these regions that were used for subsequent analyses. The goal of this model was to predict how each voxel responds to the different target objects. As an overview, we constructed a voxel-wise object encoding model using a ridge-regression approach in which beta estimates for the 80 objects were generated for each voxel and then evaluated based on how well this model predicted left out data. The natural images presented on each trial were represented as sparse 80-dimensional vectors indicating which objects were present in each image and in what quantities. We modeled the trial-wise beta estimates for each voxel using these stimulus vectors. An encoding model was created within each session, with the 6 odd runs used as training data and the 6 even runs used as test data.

#### Model estimation

Data from the 6 training runs were concatenated resulting in an 80 (items) x 375 (trials) matrix. Trial-wise betas for each voxel were regressed against the 80 item vectors using ridge-regression (Fig. 3A), which reduces overfitting by including hyperparameter λ that encourages shrinkage of model parameters. In order to determine the optimal λ for each voxel, we implemented fracridge (Rokem & Kay, 2020) in a 3-fold cross-validation procedure within the training data. In each fold, voxel betas were estimated on 66% of the training data using a range of λ values and validated on the remaining 33% of training data. On each validation trial, the model’s predicted response for each voxel was the 80-item beta estimates multiplied by the 80-item stimulus vector plus the model’s constant term. Model error was quantified as the absolute difference between the voxel’s true response on that trial and the model’s predicted response. Mean squared error (MSE) was calculated across validation trials for each λ value for each voxel, and the best λ for each voxel was extracted. For each voxel, the optimal λ values in the three iterations were averaged, and this value was used as the voxel’s λ value in the main stage of model estimation. Once the optimal λ values were extracted across voxels, the same model estimation procedure with ridge-regression was applied to the full 80 (items) x 375 (trials) training data matrix. The final output of the model estimation stage is a beta estimate for each item in each voxel, in addition to the voxel-wise constant terms.

#### Model testing

The voxel-wise beta estimates from the model estimation stage were used to predict trial-level voxel responses in the 6 left-out test runs. To generate the model’s predictions for each voxel’s response on each trial, the trial’s 80-item stimulus vector was multiplied by the voxel’s 80-item beta vector and the constant from the ridge regression was added. Each voxel’s data were mean-centered separately within the predicted and actual test patterns. Because we are interested in multivoxel object representations, we used the model’s voxel-wise predictions to generate multivoxel trial-level predictions within cubical searchlights (3 x 3 x 3 voxels) centered on each voxel across the brain. Within each searchlight, model accuracy was calculated as the Pearson correlation between predicted and actual multivoxel patterns. This correlation value was assigned to the voxel at the center of each searchlight. Thus, for each voxel in each session for each participant, we have a measure reflecting the degree to which objects are represented within its local neighborhood.

#### Generating neural representations of items

We used the encoding model to generate predicted item patterns at multiple timescales. First, in order to get the best possible model estimates, we took the optimal lambda determined in the model estimation procedure and fit the encoding model to all of the 750 trials in a given session. The beta estimates for each item were then used to generate predicted activity patterns for each of the 80 objects in each searchlight. These session-length item patterns were used in the semantic detection and semantic drift analyses described below. Second, we generated item patterns corresponding to the first and second half of each session. Each session was split into two blocks (block 1: runs 1-6; block 2: runs 7-12). Using the exact same model estimation methods described above, new encoding models were run within each block in order to generate predicted patterns (beta estimates) for each item separately within the first and second half of each session (e.g., a “giraffe” pattern in block 1, and a “giraffe” pattern in block 2). These block-length item patterns were used in the recent structure sensitivity analysis described below.

### Semantic Detection

Word2vec (Mikolov et al., 2013) was used to capture semantic relations among our 80 items. Word2vec is an implementation for word embeddings which captures similarities between words based on their co-occurrence in text. The vector embeddings for the set of items were correlated with each other to generate an 80 x 80 semantic similarity matrix. Within searchlights across the brain, we used the object encoding model to generate predicted multivoxel patterns for each of the items (at the session level) and correlated these patterns to generate an 80 x 80 neural similarity matrix (Fig. 2B). To determine whether the encoded neural patterns contained semantic information, we correlated the semantic and neural similarity matrices with each other using Spearman rank correlation (Fig. 2B). Semantically similar items tend to occur in the same images, and there is some possibility that this might lead to inflated similarity estimates between neural and semantic spaces. Thus, to ensure a conservative analytical approach, for each participant we excluded cells from the neural and semantic matrices that corresponded with items that co-occurred at any point in the experiment, resulting in a subset of item-pairs for each subject (*M*=1390 pairs, *SD*=24). The resulting Spearman’s rho coefficient was assigned to the voxel at the center of the searchlight. This analysis was run on each of the 30 sessions for each of the 8 participants. In ROI analyses, voxel values within each ROI were averaged within each session for each participant. In whole-brain analyses, brain maps were averaged across sessions.

### Long-term Drift

In order to determine whether item representations drift over long timescales, we calculated the extent to which the representation of a particular item in a particular session (e.g., giraffe in session1) is more similar to neighboring sessions (e.g., giraffe in session 2) than in distant sessions (e.g., giraffe in session 30). Within each searchlight across the brain, the encoded patterns for each item were extracted within each session and were correlated with each other to generate a 30 x 30 matrix. Cells in this matrix that corresponded to the same time lag (i.e., number of intervening sessions) were averaged, resulting in a 1 x 29 vector containing the average within-item similarities at different degrees of lag. For example, the first value in this vector would correspond to the similarity of giraffe patterns that were separated by a single session (e.g., session 1 and session 2; session 29 and session 30), whereas the last value would correspond to the similarity of giraffe patterns that were separated by 29 sessions (i.e., session 1 and session 30). This similarity-by-lag vector was regressed against the amount of lag (1-29) to determine whether within-item similarity values increased or decreased over time within each searchlight. Beta values were averaged across items, and the resulting value was assigned to the voxel at the center of the searchlight. In ROI analyses, voxel values within each ROI were averaged for each participant.

### Influence of Recent Structure

The goal of this analysis was to determine whether recent structure in the visual semantic environment—that is, the patterns of co-occurrences among items—rapidly influences the nature of those item representations.

#### Quantifying recent structure

Structure was characterized in terms of item co-occurrence across trials (Fig. 1). Co-occurrence of sets of items can be captured using the construct of network modularity or community structure, which has been used in many experimental studies of structure learning in humans (e.g., Schapiro et al., 2013; Solomon & Schapiro, 2023). In our case, the input into the modularity function is the pairwise co-occurrence frequencies of the 80 items. The modular organization of these items was calculated using Brain Connectivity Toolbox (brain-connectivity-toolbox.net; Rubinov & Sporns, 2010). The outputs of this function include an overall measure of modularity (i.e., how well the inputs can be described in terms of clusters with high within- and low between-connectivity), and assignments of each item to one of the modules detected. Thus, for each pair of items, we can observe whether the two items were assigned to the same or distinct modules (Fig. 6A). Since the modularity function is stochastic, it was run 100 times on each input; we counted, for each pair of items, the number of times (out of 100) that the items were assigned to the same module. This value was taken to reflect the measure of structural similarity between any two items. The gamma parameter in the modularity function biases the model to find smaller (γ > 1) or larger (γ < 1) modules. We set γ = 1.5 because it resulted in the most sensitivity in our structural similarity measure, as determined by a parameter search on each participant’s session 1 data. The upshot of this analysis is an 80 x 80 matrix capturing pairwise structural similarity between items, as computed by the modularity calculation (Fig. 2D).

#### Assessing influence of recent structure on neural representations

Each session was split into two blocks corresponding to the first half (runs 1-6) and second half (runs 7-12) of the session. Within each block, we calculated the co-occurrence frequencies between items across the 375 corresponding trials. These co-occurrence matrices were submitted as input to the modularity calculations described above. This enabled the quantification of structural similarity amongst items in block 1, in addition to the structural similarity amongst items in block 2. We then used our split-session encoding models to generate predicted item patterns separately for block 1 and block 2 and calculated between-item neural pattern similarity within each block. To determine whether recent structure influenced neural similarity among items, we correlated the structural similarity values from block 1 with neural similarity in block 2 (“forward correlation”). Higher correlation values indicate that items that tended to co-occur in block 1 were represented more similarity in block 2. However, this relationship on its own could indicate an influence of long-term object co-occurrence structure on neural similarity, rather than a rapid influence of recent structure. We thus also correlated structural similarity in block 2 with neural similarity in block 1 (“backwards correlation”); high correlation values indicate a relationship between long-term object co-occurrence and neural similarity, but cannot indicate a causal temporal dependence between the two. In order to control for an effect of long-term structure, we subtracted the backward correlation value from the forward correlation value. This subtraction is a powerful means of controlling for any alternative influences on neural similarity between items. Thus, positive values in this measure indicate a rapid influence of recent structure on neural representations. In ROI analyses, voxel values within each ROI were averaged within each session for each participant. In whole-brain analyses, brain maps were averaged across sessions.

### Structure-based Updating

If a relationship exists between recent semantic structure and subsequent item similarity, we aimed to determine what might modulate this effect. One possibility is that the degree to which structure modulates neural item similarity is influenced by the degree of mismatch between the recently observed structure and the long-term structure of item co-occurrences stored in memory. To make this long-term item co-occurrence structure concrete, we calculated, for each subject, the number of times each pair of items co-occurred across the 30 experimental sessions. For each session, we then generated a structure mismatch value for each session by calculating the Pearson distance between the co-occurrence statistics in the first half of the session with the long-term structure. Intuitively, this quantifies how atypical the co-occurrences were in a given session compared to the long-term average. To test whether this mismatch value relates to the degree of representational change, the relationship between structure mismatch and the recent structure effect across sessions was calculated using a Spearman correlation for each participant.

### Whole-brain significance testing

Group-level reliability across the brain was calculated using threshold-free cluster enhancement (Winkler et al., 2014) on the session-averaged maps from each participant; significance was assessed at a corrected α = 0.05. The resulting maps were used to create masks within which subsequent reliability analyses were performed; to increase power for these subsequent analyses (by reducing multiple comparisons), masks were generated using a threshold of α = 0.01. The first step was to locate regions of the brain in which object encoding was successful. Within the resulting encoding mask, regions that reliably contained semantic content were located. Within the resulting semantic mask, the reliability of long-term drift effects and short-term structural effects were assessed.

## Acknowledgements

We are grateful to Michael Arcaro for helpful discussions. Collection of the NSD dataset was supported by NSF IIS-1822683 and NSF IIS-1822929.

