## Supplemental Material for "Semantic plasticity across timescales in the human brain"

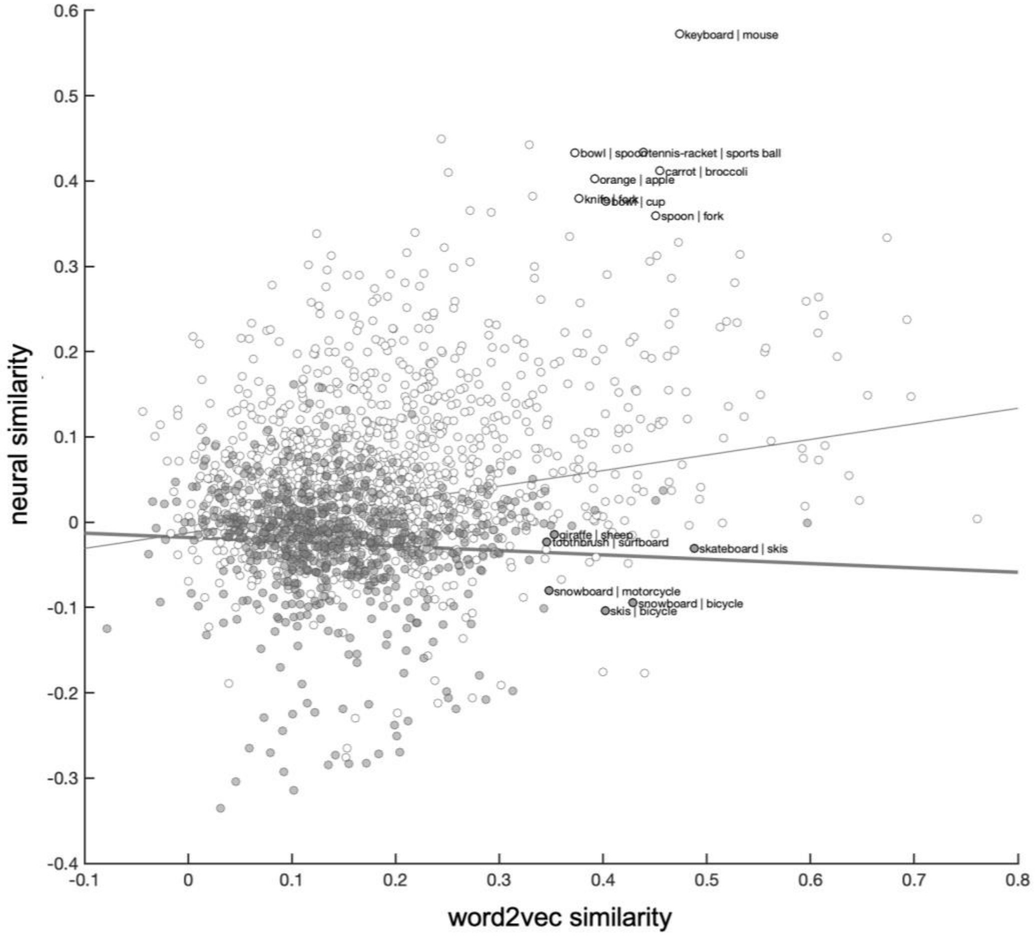


**Supplemental Figure 1.** Visualizing the relationship between word2vec similarity and neural similarity across items in V1, averaged across participants. We used a conservative approach in localizing semantic regions in order to ensure that similarity between items could not be driven by their co-occurrence in the visual stimuli: In examining the relationship between word2vec and neural similarity, we excluded pairs of items that co-occurred at any point throughout the experiment (white circles), leaving only items that did not co-occur (grey circles). When all item pairs are included in the analysis, a positive relationship between word2vec and neural similarity is observed in V1 (thin trend line). However, when only considering items that did not co-occur, this relationship is negative (thick trend line). This trend is driven by item pairs with high word2vec similarity and low neural similarity: these pairs tend to include items that are highly conceptually similar, but would never be found in the same visual environment. For example, one will find skis on mountain slopes and skateboards in skate parks, but never vice versa. Because V1 is sensitive to the visual environment, if two items always appear in dissimilar environments (e.g., mountains vs. skate parks), the representation of those items in V1 will also be dissimilar. On the other hand, it is possible for two dissimilar items to never co-occur yet appear in the same environment (e.g., cat and fork). Thus, the negative correlation between word2vec and neural similarity in V1 is a consequence of our exclusion of co-occurring items in our semantic analysis.

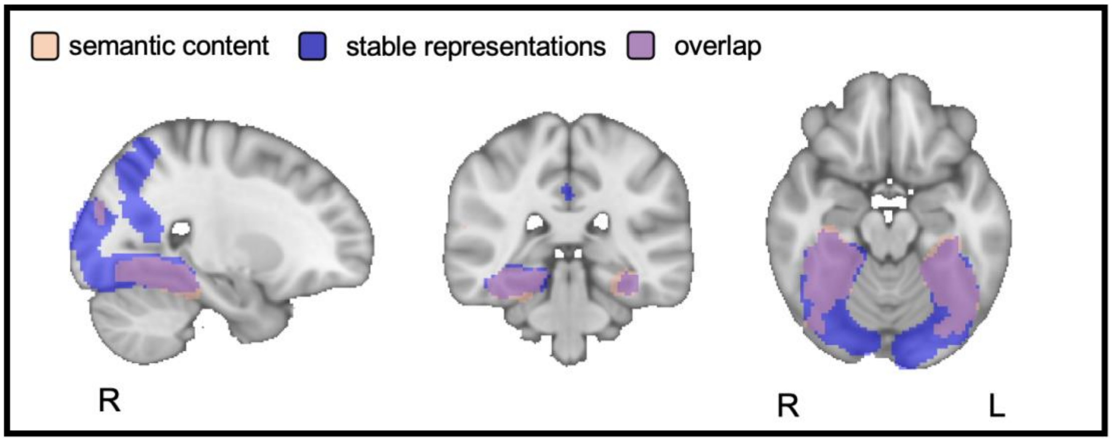


**Supplemental Figure 2.** Based on the group-level drift results reported in Fig. 5C, it was theoretically possible that the more posterior semantic regions did not exhibit representational drift not because the representations therein were more stable, but because there was no stability between sessions at all, which would preclude the ability for us to detect drift. For example, if there is no correspondence between an item's representation in session 1 and session 2, there would be no basis for detecting a decrease in that correspondence over time. In order to rule out this possibility, we analyzed the short-term stability of item representations across the brain by assessing whether each item's representation positively correlated with itself in neighboring sessions (e.g., sessions 1 and 2; sessions 29 and 30). Within each voxel, for each participant, these neighboring session similarities were combined within and then across items. Group-level significance was assessed within the same encoding mask used in previous analyses. Regions exhibiting short-term representational stability are shown in blue; the semantic regions reported in the main manuscript are shown in pink (Fig. 4); overlap between these regions is shown in purple. Almost all semantic regions exhibit short-term representational stability. This analysis demonstrates that the lack of a drift effect in more posterior semantic regions (Fig. 5C) cannot be explained by a complete lack of representational stability in those regions. Instead, regions that revealed significant levels of drift can be interpreted to exhibit increased plasticity relative to the semantic regions that did not.

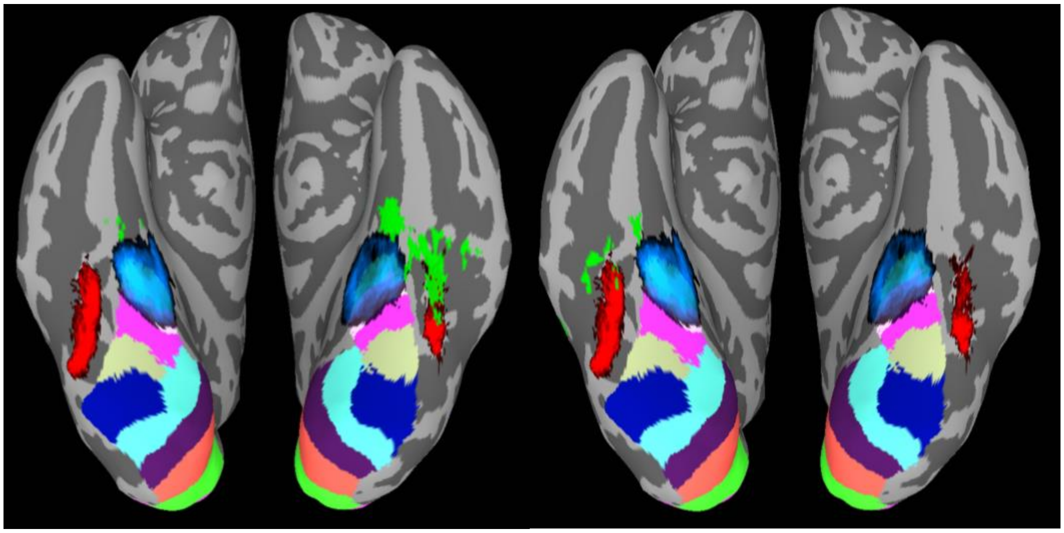

**Supplemental Figure 3.** Visualizing the location of the short- and long-term plasticity effects on the ventral cortical surface relative to parahippocampal place area (PPA), fusiform face area (FFA), and retinotopic visual areas, displayed on an MNI template brain. Solid color areas correspond to the ventral regions of the Wang retinotopic atlas (Wang et al., 2015). The blue and red graded areas correspond to probabilistic maps of PPA and FFA, respectively, with lighter colors corresponding to higher probabilities (Arcaro et al., 2018). Left: Regions showing long term representational drift overlaid in lime green. Right: Regions showing sensitivity to recent statistical structure overlaid in lime green.
